# Investigating the Role of Neck Muscle Activation and Neck Damping Characteristics in Brain Injury Mechanism

**DOI:** 10.1101/2023.11.15.567289

**Authors:** Hossein Bahreinizad, Suman K. Chowdhury

**Affiliations:** Department of Industrial, Manufacturing and Systems Engineering, Texas Tech University, Lubbock, Texas, USA

**Author notes:** All correspondence should be addressed to—Dr. Suman K. Chowdhury, Assistant Professor, Department of Industrial, Manufacturing and Systems Engineering, Texas Tech University 905 Canton Ave, Lubbock 79409-3061, Texas, USA.

**Keywords:** Simulation and modeling, Finite element method, Traumatic brain injury, Computational biomechanics, Neck damping, Active muscles

## Abstract

**Purpose:** This study aimed to investigate the role of neck muscle activity and neck damping characteristics in traumatic brain injury (TBI) mechanisms.

**Methods:** We used a previously validated head-neck finite element (FE) model that incorporates various components such as scalp, skull, cerebrospinal fluid, brain, muscles, ligaments, cervical vertebrae, and intervertebral discs. Impact scenarios included a Golf ball impact, NBDL linear acceleration, and Zhang’s linear and rotational accelerations. Three muscle activation strategies (no-activation, low-to-medium, and high activation levels) and two neck damping levels by perturbing intervertebral disc properties (high: hyper-viscoelastic and low: hyper-elastic) strategies were examined. We employed Head Injury Criterion (HIC), Brain Injury Criterion (BrIC), and maximum principal strain (MPS) as TBI measures.

**Results:** Increased neck muscle activation consistently reduced the values of all TBI measures in Golf ball impact (HIC: 4%-7%, BrIC: 11%-25%, and MPS (occipital): 27%-50%) and NBDL study (HIC: 64%-69%, BrIC: 3%-9%, and MPS (occipital): 6%-19%) simulations. In Zhang’s study, TBI metric values decreased with the increased muscle activation from no-activation to low-to-medium (HIC: 74%-83%, BrIC: 27%-27%, and MPS (occipital): 60%-90%) and then drastically increased with further increases to the high activation level (HIC: 288%-507%, BrIC: 1%-25%, and MPS (occipital): 23%-305%). Neck damping changes from low to high decreased all values of TBI metrics, particularly in Zhang’s study (up to 40% reductions).

**Conclusion:** Our results underscore the pivotal role of neck muscle activation and neck damping in TBI mitigation and holds promise to advance effective TBI prevention and protection strategies for diverse applications.

## Introduction

Traumatic brain injury (TBI) occurs when a sudden mechanical impact to the head or body leads to structural and functional changes in the brain [1]. According to the Global Burden of Diseases, Injuries, and Risk Factors (GBD) report, the prevalence of TBI increased by 24.4% from 1990 to 2019, with an estimated 49 million cases globally in 2019. In the United States alone, the Centers for Disease Control and Prevention Wonder database reported 69,473 TBI-related deaths in 2021 and 223,135 TBI-related hospitalizations in 2019 [2]. The leading causes of TBI are falls, followed by motor vehicle accidents, assault, and firearm-related injuries [3–5]. Sports (e.g., football, hockey, etc.) and recreational activities (e.g., bicycling, horse riding, and playground activities) are also major causes of TBIs [6, 7]. According to a 2019 CDC report approximately 283,000 children (under the age of 18) visited the emergency department due to TBIs associated with sports and recreational activities [8]. Previous TBI studies have identified the unrestricted, rapid acceleration of the head as the main factor contributing to TBIs [9, 10]. This rapid head motion, primarily governed by the action of neck muscles, is also influenced by the neck damping characteristics, which stem primarily from the cervical intervertebral discs and ligaments [11, 12]. Thus, in order to develop effective preventive measures, it is crucial to investigate how neck muscles and neck damping characteristics influence the TBI mechanism, i.e., the rapid deformation and strain of the brain.

Previous literature showed a range of experimental (both in-vivo and in-vitro studies) and computational methods to study the TBI mechanism. In-vivo experimental studies primarily utilized quantitative video analysis [13] and wearable sensors [14] to understand the injury characteristics and TBI mechanisms based on head kinematics. However, these approaches are limited in their ability to measure internal parameters of the brain and other head components. On the other hand, in-vitro experimental studies used anthropometric test device [15] and/or cadavers [16], to explore the mechanical response of internal head components. Nevertheless, both experimental approaches may not fully capture the complexities of the living human body. To overcome these limitations of experimental methods, researchers have turned to computational approaches in TBI research. Notably, finite element (FE) models have gained popularity as powerful tools for studying TBI. These models can offer a highly detailed representations of the biomechanical behavior of head and neck structures, enabling the study of both external kinematics and internal responses of the brain and other head components during various impact scenarios [11, 12, 17, 18]. Thus, by leveraging FE-based head-neck computational models, researchers can overcome the limitations of experimental methods and gain a more comprehensive understanding of TBI mechanics and associated preventive measures.

In general, cervical intervertebral discs acts as damper in the head-neck system, thereby absorbing the impact forces and reducing the risk of brain and other head-neck injuries [19]. Furthermore, the surrounding neck muscles play a crucial role in counteracting the external forces and mitigating the injury severity of the head-neck system [20]. Neck muscles also exert compressive forces to counteract the tensile force induced by the external impact forces acting on the head-neck system. The direction and magnitude of these compressive forces depend on individual neck muscles’ maximum strength, activation level, position, and their roles a flexor, extensor, or rotator muscle [21]. Accordingly, some experimental studies investigated the role of neck muscles in reducing the head-neck injury severity, however, they reported conflicting findings – Some studies [22, 23] found a significant reduction in TBI risk associated with increased neck muscle strength, whereas other studies [24–26] reported no significant effect. To address these putative results, Jin et al. [11] employed a computational head-neck model to investigate the impact of neck muscle activation levels and strategies on brain injury mechanism. They observed that though an increased muscle activation level reduces the brain injury risk, the combined muscle activation strategies had a more significant effect on it. In another head-neck FE study by Bruneau et al. [17], the increased muscle activation was found to cause reduced head-neck kinematics. However, both these studies [11, 17] did not consider the influence of neck damping characteristics. On the contrary, Dirisila et al. [12] investigated the influence of neck damping on the brain response to a direct frontal impact to the head using a rigid impactor and modeled the neck as a simple spring-damper system (instead of modeling biofidelic cervical vertebrae, muscles, intervertebral discs, and ligaments) in their head-neck FE model. Their observations indicated that neck damping, characterized by viscoelastic properties, had minimal effects on the brain response in the initial stages (simulation time) of an impact but subsequently reduced the brain’s stress response in the later stages of the impact. Findings of these studies highlight the necessity of further research on the development of a FE model incorporating accurate neck muscle activation strategies and damping properties of intervertebral discs, in addition to explore how neck muscle activation and neck damping properties can collectively affect the TBI mechanism.

Therefore, the goal of this study was to explore the role of neck muscle activity and neck damping characteristics in TBI mechanism. Specifically, we aimed to assess the effects of various muscle activation strategies (no activation, low/medium activation, and high activation) and neck damping properties (hyper-viscoelastic/hyper-elastic intervertebral disks) on the mechanical response of the head-neck system under various impact scenarios, including Golf ball impacts, linear acceleration, and combined linear and rotational acceleration. By conducting these simulations, we seek to gain a comprehensive understanding of the mechanisms underlying TBI protection and provide insights for the development of effective preventive strategies. This research has the potential to enhance our knowledge of the neck’s role in mitigating brain injuries and contribute to improved safety measures in impact-related activities.

## Materials and Methods

### Head-neck model

We used our previously validated head-neck FE model to investigate our study objective [27]. The model incorporates various components, including the scalp, skull, cerebrospinal fluid (CSF), brain, dura mater, pia mater, neck muscles, neck ligaments, cervical vertebrae (C1-C7), and intervertebral discs [27]. This model was created using head and neck MRI data derived from a male participant (age: 42 years, height: 176 cm, and weight: 106 kg), resulting in a comprehensive structure comprising 1.36 million elements. In this model, the scalp, skull, and neck segments contributed 0.27 million, 0.23 million, and 0.072 million tetrahedral elements, respectively. Additionally, the model featured 0.1 million quad shell elements, with the dura and pia maters accounting for 0.05 million and 0.04 million elements, respectively. We modeled the skull, pia mater, dura mater, and vertebrae as linear elastic materials. The scalp was represented using a linear viscoelastic material, the CSF as a hyper-elastic material, the intervertebral discs as hyper-viscoelastic materials, and the brain’s gray and white matter as a hyper-viscoelastic material. One notable feature of this model is its incorporation of the Hill-type muscle model [28], allowing us to simulate active muscle behavior and investigate the impacts of muscle activation on our study objectives. Moreover, we modeled the intervertebral discs with a hyper-viscoelastic material model [29]. The material properties of individual head-neck structures are provided in Table 1 [27].

**Table 1.**
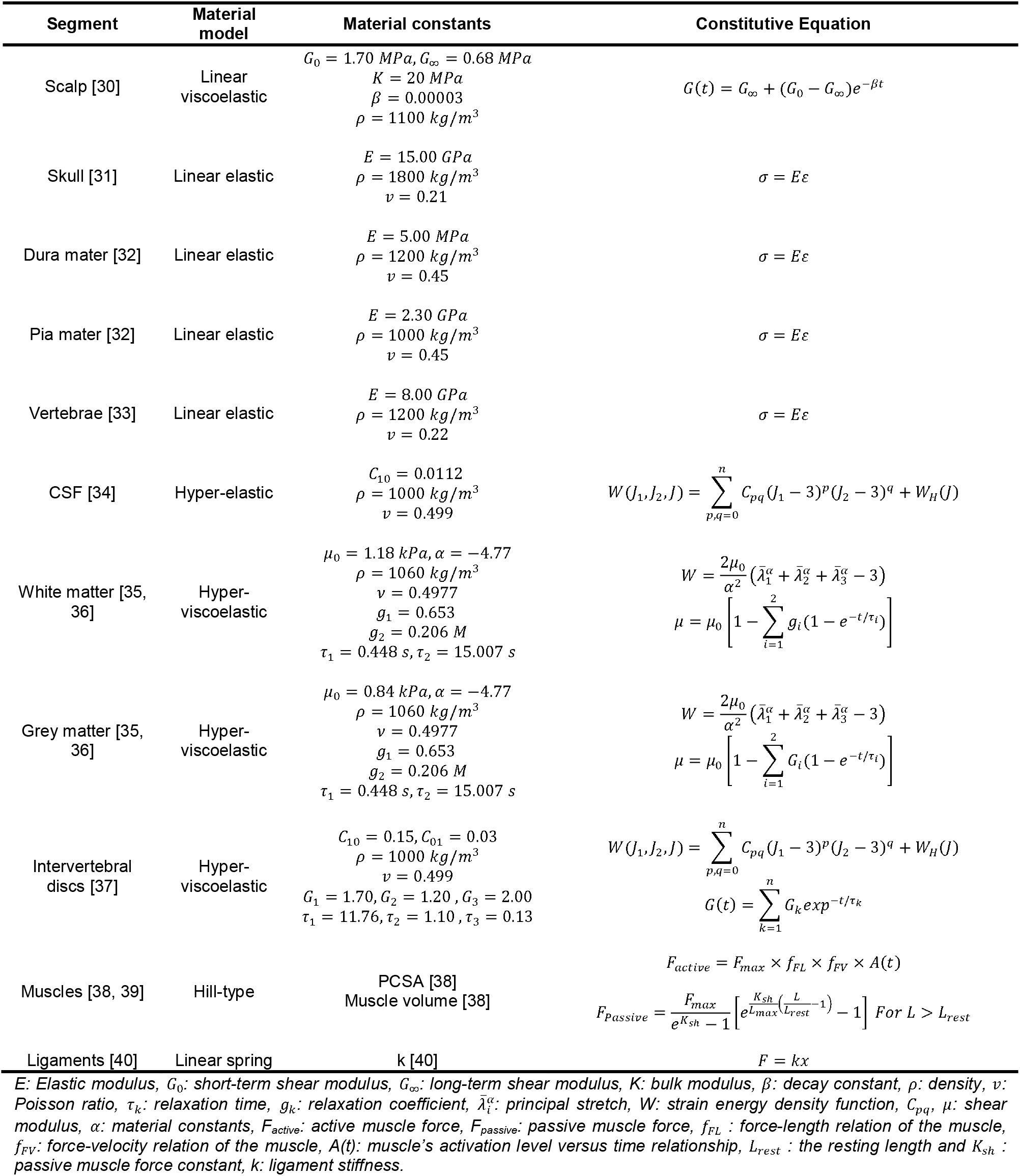
Material properties and constitutive equations of each head-neck structure included in our model.

### Impact Loads

We simulated three impact conditions (Fig. 1). To address the roles played by the neck under focal injury conditions (confined to brain damage at one specific area), we simulated a Golf ball impact scenario, as outlined in a previous study [41]. Our simulation included a Golf ball hitting the frontal area of the head model with an initial velocity of 35 m/s. The Golf ball was represented as a three-layered structure consisting of an inner core, a mantle, and a cover. The core and mantle were modeled using hyper-viscoelastic materials, while the cover was modeled using a hyper-viscoelastic material (Table 2) [3]. Equation 1 describes the hyper-elastic behavior of Golf ball layers, whereas Equation 2 describes their viscoelastic response.

**Fig. 1.**
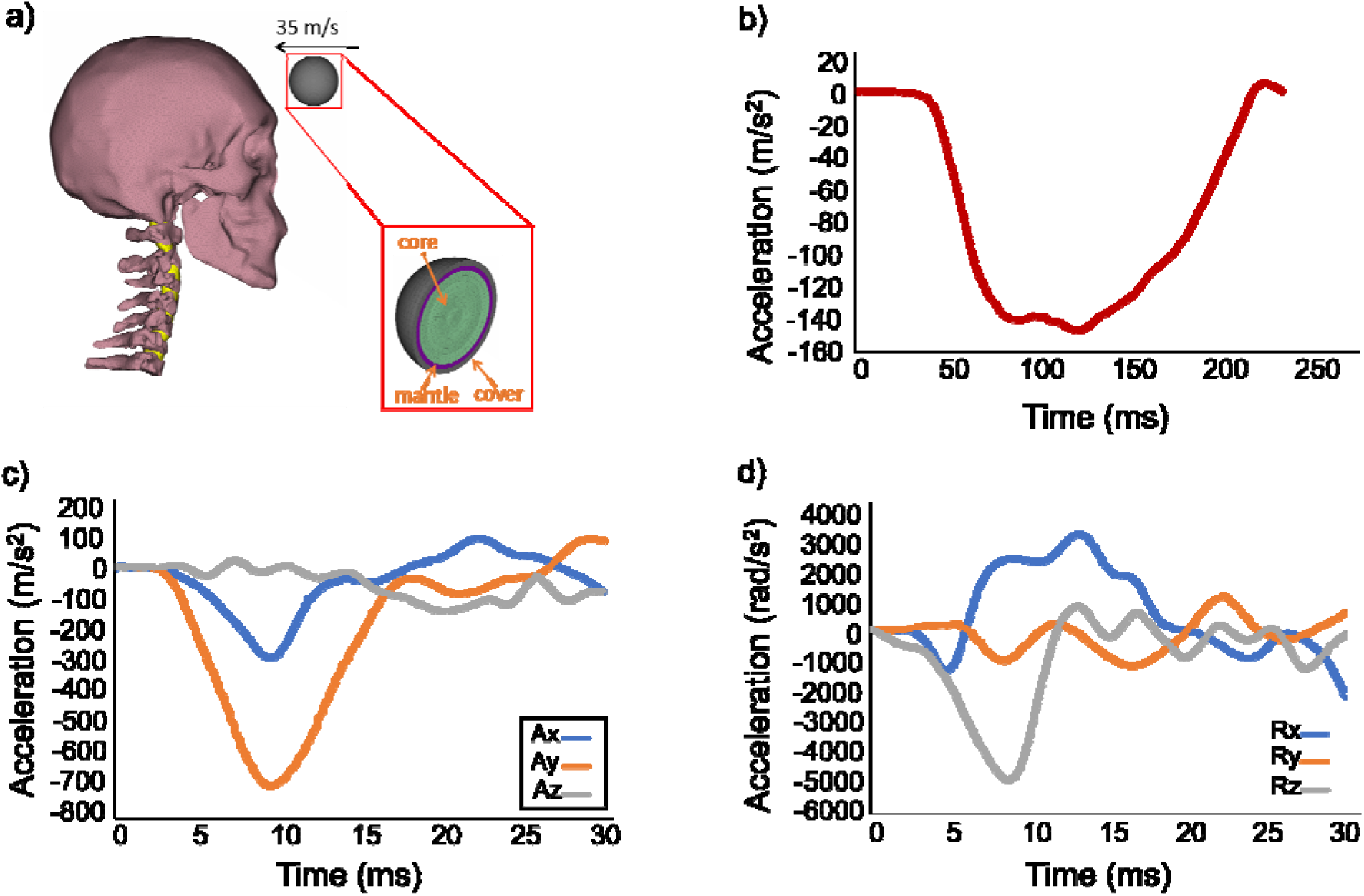
Impact loads simulated in this study include: (a) Golf ball impact simulation (muscles, ligaments, and scalp are excluded from the figure for better visualization) [41], (b) Naval Biodynamics Laboratory (NBDL) study’s linear acceleration profile [42], (c) Linear acceleration (Ax, Ay, Az) profiles, and (d) Rotational acceleration (Rx, Ry, Rz) profiles of helmet-to-helmet collisions in American football, taken from Zhang’s study [43].

**Table 2.** Mechanical characteristics of the Golf ball [3].

| | Outer diameter (mm) | No. of elements | $\rho$ (kg/m<sup>3</sup>) | $\nu$ | $C_{01}$ (MPa) | $C_{10}$ (MPa) | $G_1$ (MPa) | $\beta_1$ (S<sup>-1</sup>) | $G$ (MPa) | SIGF (kPa) |
| --- | --- | --- | --- | --- | --- | --- | --- | --- | --- | --- |
| Core | 35.4 | 8580 | 1125 | 0.49 | 1.50 | 6.00 | 10.50 | 37000 | 4.50 | 9.00 |
| Mantle | 38.8 | 1800 | 1274 | 0.49 | 3.67 | 14.70 | 18.30 | 18000 | 18.30 | 36.70 |
| Cover | 42.8 | 2040 | 950 | 0.49 | 10.50 | 42.00 | - | - | - | - |

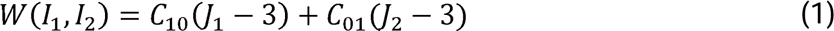

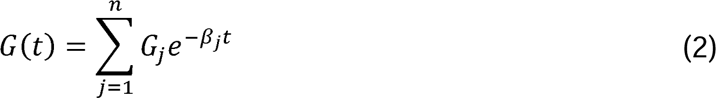

In Equations 1 and 2, *J* is invariant of the right Cauchy-Green deformation tensor, W is the strain energy density function, is the decay constant, G_j_ is the shear relaxation modulus, G is shear modulus for frequency independent damping and SIGF is the stress limit for frequency independent damping. We connected the Golf ball’s three layers using a tied boundary condition.

Our second and third impact scenarios were carefully selected to investigate the role of neck in diffuse injuries, which involve more extensive disruption of brain tissue and can affect multiple areas simultaneously [9, 10, 44]. As diffuse injuries are primarily associated with acceleration scenarios [10], we incorporated two acceleration scenarios: 1) the linear acceleration scenario from the Naval Biodynamics Laboratory (NBDL) study [45] and 2) the combined linear and rotational acceleration scenario proposed by Zhang et al. [43]. The NBDL acceleration profile was applied to the entire head-neck model, while the Zhang’s acceleration profile specifically targeted the center of mass of the head. The Zhang’s acceleration scenario was derived from the recreation of American football accidents. Considering the ongoing debate surrounding the main cause of injuries, whether it is linear or rotational acceleration, it was imperative for us to include both scenarios in our study.

### Study design

In order to examine the effects of neck muscle activations and neck damping characteristics on mechanical responses of the brain and the head-neck system, we considered three activation strategies across all neck muscles: 1) no-activation level, representing unprepared muscles experiencing an unanticipated impact scenario, 2) a low-to-medium activation level, indicating weak or inadequate muscle activation, and 3) high activation level, simulating tensed muscle conditions under anticipated impact scenarios. As previous research indicated that spinal muscles exhibit slight activity (approximately 5%) to provide stability to the spine (even when they are not actively contracting to provide any mobility), we applied a 5% activation level to all neck muscles during the no-activation level [46]. For second and third activation strategies, we provided varied activation levels to neck flexor and extensor muscle groups in order to simulate the realistic kinematic response of the head and neck system. In the Golf ball impact study, the flexor muscles play a crucial role in counteracting the frontal impact force. Therefore, for the low-to-medium activation level, we applied a 25% activation level to the flexor muscles and 10% to the extensor muscles. Similarly, for the high activation level, we simulated tensed muscle conditions by using 80% flexor muscle activation and 10% extensor muscle activation.

In NBDL acceleration simulations, we applied a 25% activation level to extensor muscles and 10% to flexor muscles for the low-to-medium activation level as neck extensors are generally activated more than neck flexors in order to attenuate neck hyper-flexion. For the high activation level, we increased the extensor muscle activation to 80% while maintained the flexor muscle activation at 10%. In Zhang’s combined linear and rotational acceleration scenarios, which do not strictly fall under pure neck flexion or extension, we applied same activation level to all neck flexor and extensor muscles throughout the impact duration. Specifically, we applied 25% activation level during the low-to-medium activation simulations and 80% during the high activation level simulation across all neck flexors and extensors.

The influence of neck damping on the risk of brain injury was examined by modeling the cervical intervertebral discs as hyper-elastic (low neck damping) and hyper-viscoelastic (high neck damping) properties. The hyper-elastic properties includes a nonlinear elastic module characterized by Mooney Rivlin hyper-elastic formulation [47, 48], whereas the hyper-viscoelastic properties include both the hyper-elastic component and a Prony series component to account for viscoelastic properties of the intervertebral discs [37]. We examined the effects of neck damping on the TBI mechanism by switching between hyper-elastic and hyper-viscoelastic material models of the intervertebral discs.

### Injury estimation

Over time, researchers have made significant efforts to establish various injury criteria that can reliably predict the occurrence of brain injury. Initially, linear kinematics of the head were used as primary parameters in popular injury criteria like Head Injury Criterion (HIC) [49] and Gadd Severity Index (GSI) [50]. Later, it was found that the mechanical tolerance of brain tissues are poor against rotational acceleration and thus suggested to consider head rotational kinematics as another primary parameter in injury criterial formulation [51]. Consequently, alternative head injury criteria based on rotational kinematics were developed. For example, Takhounts et al. [51] devised brain injury criteria (BrIC) by correlating head rotational kinematics with brain strain. While the ease of measuring head kinematics made them a practical and popular option, the measurement of brain tissue deformation, such as maximal principal strain (MPS), has been proven to be more accurate and reliable measure of predicting brain injury, in particular, in diffuse brain injury scenarios [52, 53]. However, the measurement of brain MPS requires FE-based computational models, and is infeasible to assess in experimental studies [14]. As a result, the HIC [49] has been widely used as a TBI predictive metric (Equation 3) in both experimental and computational studies.

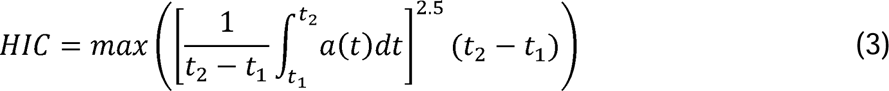

The variable, t represents time in milliseconds (ms), while a(t) denotes the linear acceleration profile measured in g’s. The HIC quantifies the maximum value of the average acceleration over a specific time duration, raised to the power of 2.5 and multiplied by the duration itself. It is recommended to impose a limit on the time duration to enhance the relevance of HIC. In this study, we adopted HIC_15_, which restricts the time difference between t_2_ and t_1_ to a maximum of 15 ms. HIC_15_ is a widely implemented variant of HIC, known for its popularity and acceptance in the field.

As HIC does not consider the rotational kinematics of the head, the BrIC was also widely used [51] as an additional parameter for estimating brain injuries (Equation 4).

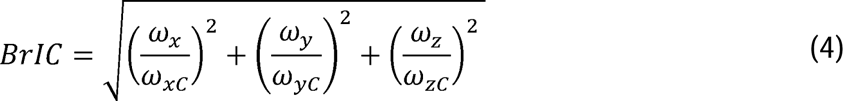

In Equation 5, *ω_x_*, *ω_y_*, and *ω_z_* denotes angular velocity in x, y, and z directions respectively. Conversely, *ω_xc_*, *ω_yc_*, and *ω_zc_* stands for the critical value of these parameters. We adopted these critical values from the literature with values of 66.25, 56.45, and 42.87 rad/s for *ω_xc_*, *ω_yc_*, and *ω_zc_* respectively [51]. Using these BrIC values, we can estimate the abbreviated injury scale (AIS) spanning from 1 to 5 [51].

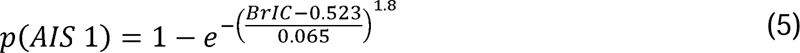

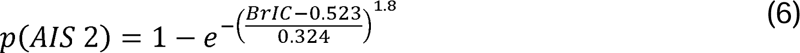

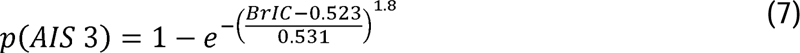

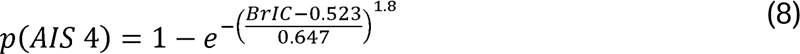

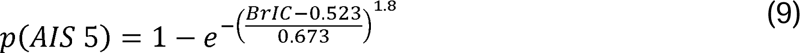

Equations 5 – 9 indicate the formulations for estimating the probability of AIS1, AIS2, AIS3, AIS4, and AIS5 scales that respectively indicate minor, moderate, serious (not life-threatening), severe (life-threatening), and critical (survival uncertain) severity of brain injuries. Furthermore, as our biofidilic head-neck FE model allowed us to go beyond the aforementioned kinematics-based injury metrics, we computed the brain’s MPS in frontal, parietal, and occipital regions, alongside HIC and BrIC measures. We used the LS-DYNA (Livermore Software Technology Corporation, USA) explicit solver for conducting all impact simulations. These simulations were executed on the high-performance computing center located at our university. The computing center boasts cutting-edge infrastructure, including two AMD EPYC 7702 CPUs and 500 GB of memory per node, which provided ample computational resources to meet the demands of our simulations. Following the completion of the simulations, we performed the postprocessing of the results using META (BETA CAE Systems SA, Greece) and MATLAB R2021b (MathWorks, USA). This combination of software tools allowed us to effectively analyze and interpret the obtained data.

## Results

### Analysis of HIC and BrIC Values

In both Golf ball impact and NBDL study simulations, the increase in muscle activation consistently led to a decrease in HIC and BrIC (Fig. 2) values. In comparison to no-activation baseline, Golf ball impact simulation showed a decrease in in both HIC and BrIC values for both low-to-medium activation (HIC: 4% for both low and high damping; BrIC: 13% for high damping, and 11% for low neck damping) and high activation level (HIC: 7% for both low and high damping; BrIC: 25% for both low and high damping). In NBDL simulations, while compared to the no-activation level, the reductions in HIC values were more substantial for both low-to-medium (HIC: 64% for high damping, and 67% for low neck damping; BrIC: 3% for high damping, and 5% for low neck damping) and high activation levels (HIC: 67% for high damping, and 69% for low neck damping; BrIC: 8% for high damping, and 9% for low neck damping). On the contrary, both HIC and BrIC values exhibited interesting trends in Zhang’s study simulations. At first, the change in muscle activation from no-activation to low-to-medium levels decreased the HIC values by about 83% (high neck damping) and 74% (low neck damping) and the BrIC values by about 27% (regardless of the neck damping). Then, a drastic increase in HIC values (about 507% with high and 288% with low neck damping) and a moderate increase in BrIC values (about 25% with high and 1% with low neck damping) were observed with the increased muscle activation from low-to-medium to high activation levels.

**Fig. 2.**
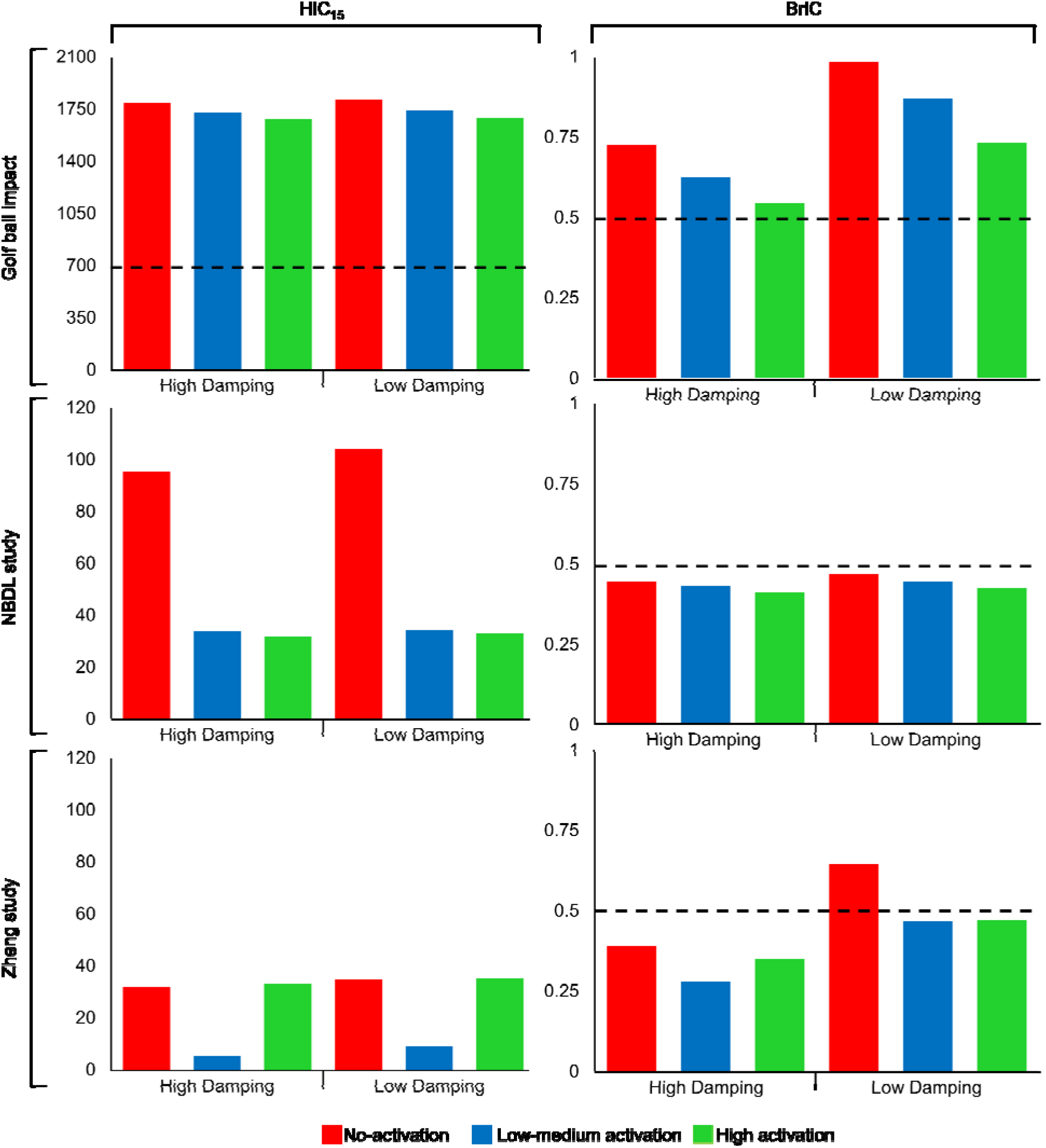
Comparative analysis of HIC_15_ and BrIC in three simulated scenarios—Golf ball impact, NBDL’s linear acceleration, and Zhang’s combined rotational and linear acceleration simulations—for two neck damping—low (hyper-elastic) and high (hyper-viscoelastic)—properties and three muscle activation levels: 1) no-activation (unprepared muscles), 2) low-to-medium activation (weak or inadequate muscle activation), and 3) high activation (simulating tensed muscles). The dashed line indicates the injury threshold level.

The change in neck damping from low to high consistently showed decreasing trends of HIC and BrIC (Fig. 2) values across all simulations. In particular, Zhang’s study simulations exhibited a substantial impact of neck damping (about 9%, 40%, and 6% reductions in HIC values and about 39%, 40%, and 25% reductions in BrIC values for no-activation, low-to-medium, and high activation levels, respectively). The NBDL study simulations revealed moderate impacts of neck damping – approximately 8%, 1%, and 3% reductions in HIC values and 5%, 3%, and 4% reductions in BrIC values for no-activation, low-to-medium, and high muscle activation levels, respectively. In contrast, the effect of neck damping in Golf ball impact simulations showed a trivial change in HIC values (about 1% reduction) but a moderate change in BrIC values (about 26% reduction) across all muscle activation levels.

### Analysis of Brain MPS Results

In both Golf ball impact and NBDL study (Fig. 3) simulations, the brain MPS in frontal, parietal, and occipital regions consistently decreased with an increase in muscle activation level and neck damping. The most notable reductions were observed in the occipital region – about 17% (high neck damping) and 6% (low neck damping) for low-to-medium activation level and 18% (high neck damping) and 19% (low neck damping) for high activation level for Golf ball impact simulations and approximately 27% (high neck damping) and 29% (low neck damping) for low-to-medium activation level and 50% (high neck damping) and 41% (low neck damping) for high activation level in NBDL study simulations, compared to the no-activation level.

**Fig. 3.**
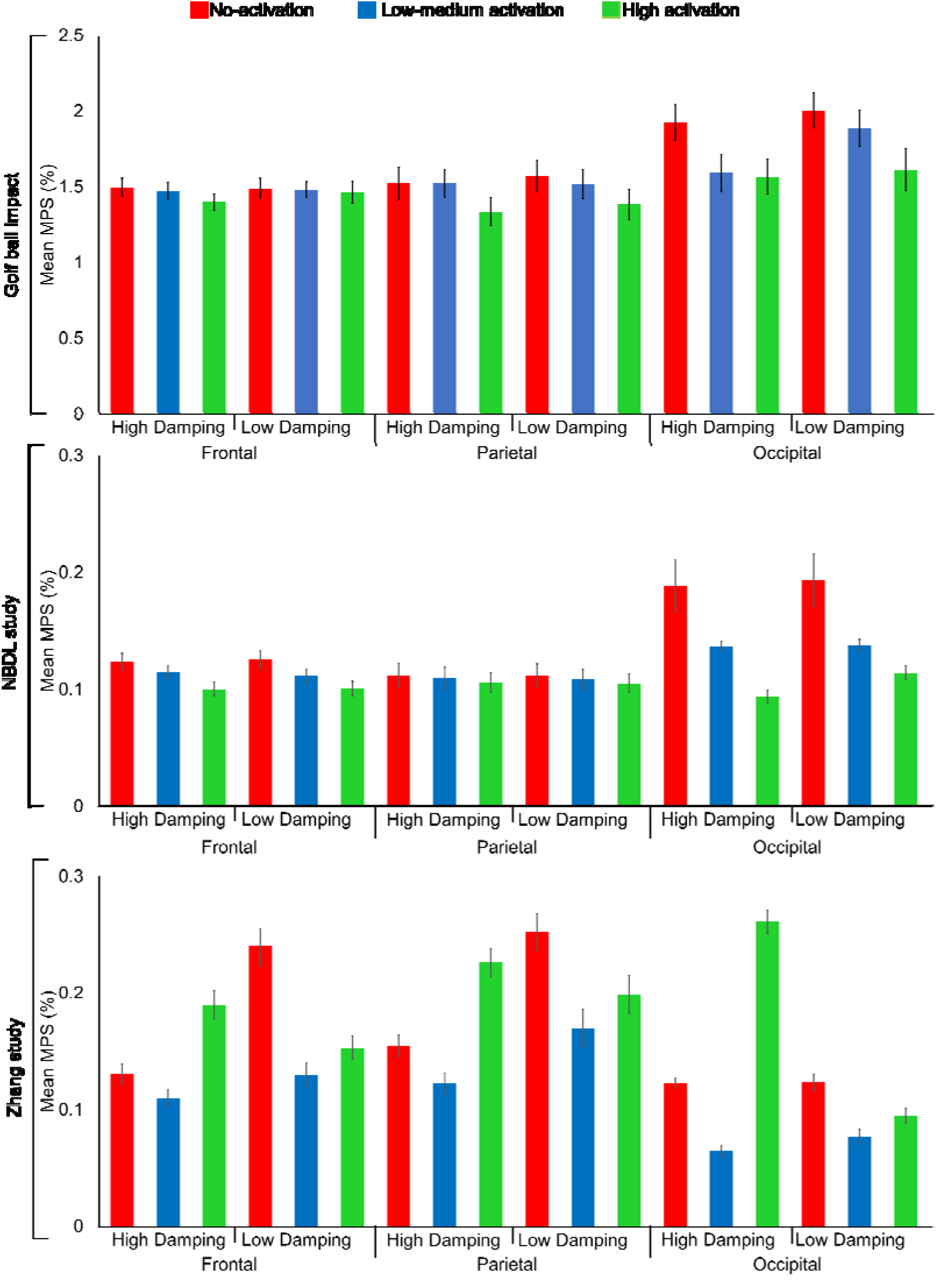
Comparative analysis of the brain maximum principle strain (MPS) (mean ± standard error) at three frontal, parietal, and occipital regions under three muscle activation levels—1) no-activation (unprepared muscles), 2) low-to-medium activation (weak or inadequate muscle activation), and 3) high activation (simulating tensed muscles)—and for two neck damping—low (hyper-elastic) and high (hyper-viscoelastic)—properties in three simulated scenarios: Golf ball impact, NBDL’s linear acceleration, and Zhang’s combined rotational and linear acceleration simulations.

In Zhang’s study simulations, similar to HIC and BrIC results, the brain MPS in all three regions notably decreased with the change in muscle activation from the no-activation to low-to-medium activation levels, but increased from the low-to-medium to high activation levels (Fig. 3). In the frontal region, we observed a decrease of about 19% (high neck damping) and 84% (low neck damping) at low-to-medium activation level compared to no-activation level and, conversely, an increase of about 73% (high neck damping) and 18% (low neck damping) at high activation level than the low-to-medium activation levels. In the parietal region, there was a decrease of about 26% with high and 48% with low neck damping at low-to-medium activation level compared to no-activation level an increase of about 84% with high and 17% with low neck damping at high activation level than the low-to-medium activation level. In the occipital region, the increased muscle activity from no-activation to low-to-medium activation levels reduced the brain MPS by about 90% (high damping) and 60% (low neck damping) and increased by about 305% (high damping) and 23% (low damping) with from the low-to-medium to high activation levels. The increase in neck damping reduced the brain MPS by approximately 83%, 28%, and 1% at no-activation and increased them by about 18%, 38%, and 19% at low-to-medium, along with around 19%, 12%, and 63% at high activation levels at frontal, parietal, and occipital regions, respectively.

### Probability of TBI

In NBDL and Zhang’s study simulations, the HIC and BrIC values remained below the injury threshold, except for one scenario (no-activation level, low neck damping) in Zhang’s study wherein the probable risks of AIS1, AIS2, AIS3, AIS4, and AIS5 based on BrIC values were about 98.54%, 20.71%, 9.11%, 6.45%, and 6.04%, respectively. On the contrary, both HIC and BrIC values showed the risk of TBIs in all scenarios of Golf ball impact study (Fig. 3). Therefore, we only estimated the AIS 1 – 5 for the Golf ball impact simulations. As shown in Fig. 4, the AIS values of Golf ball impact simulations showed a trend of reduced probability of TBI with the increase in muscle activation. This reduction was particularly large with higher neck damping (about 4% for AIS 1 and 60% for AIS 2-5 at the low-to-medium activation level and about 69% for AIS 1 and 96% for AIS 2-5 at the high activation level) than the lower neck damping (about 1% for AIS 1, 16% for AIS 2, and roughly 30% for AIS 3-5 at the low-to-medium activation level and about 1% for AIS 1, 50% for AIS 2, and 65% for AIS 3-5 for the high activation level).

**Fig. 4.**
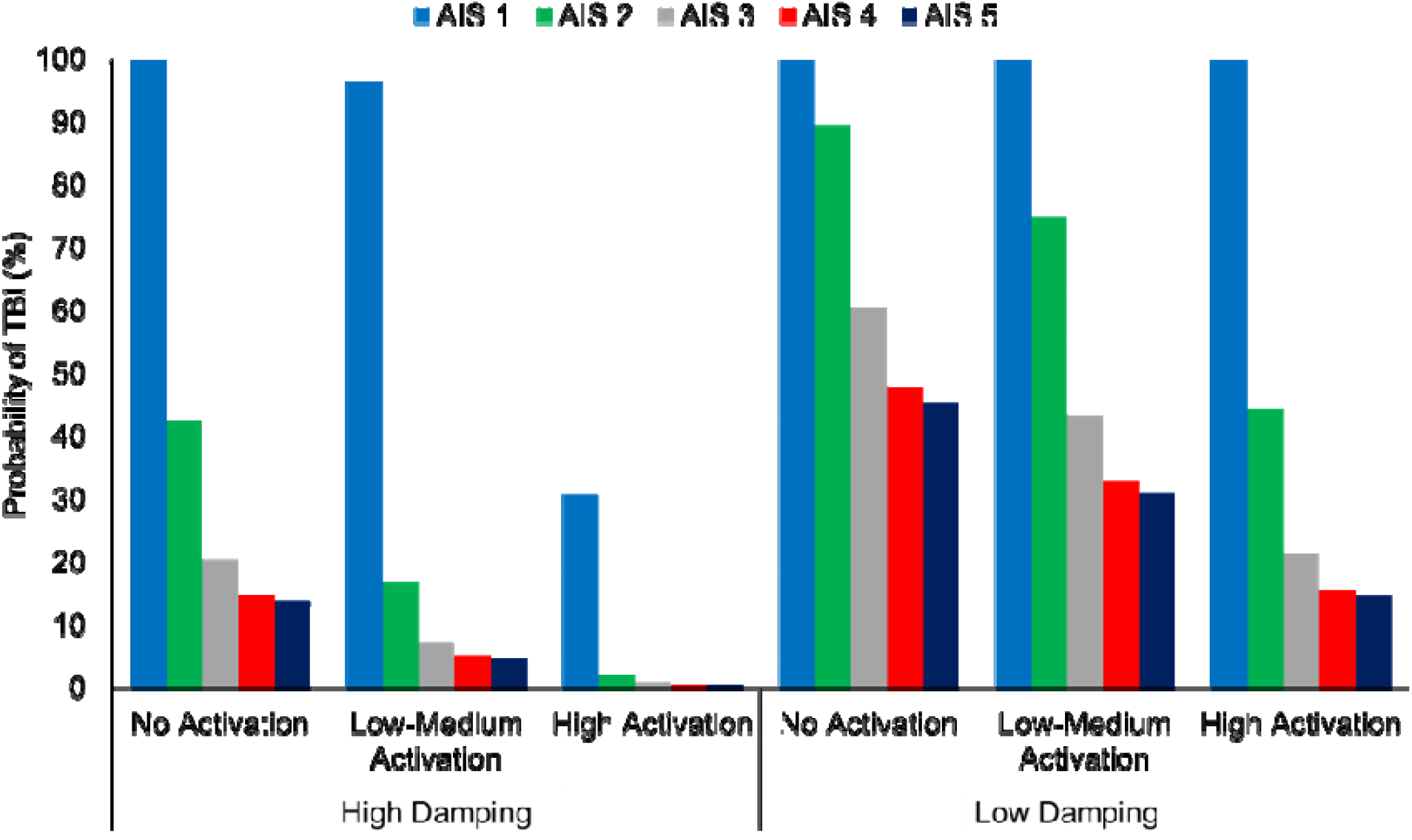
Probability of traumatic brain injury (TBI) assessed using abbreviated injury scale (AIS) for the Golf ball impact simulations. AIS1, AIS2, AIS3, AIS4, and AIS5 indicate minor, moderate, serious (not life-threatening), severe (life-threatening), and critical (survival uncertain) severity of injuries, respectively.

## Discussion

We aimed to study the impacts of neck muscle activation and neck damping on TBI outcomes for three impact scenarios—1) direct impact (Golf ball impact study), 2) pure linear acceleration (NBDL study), and 3) combined linear and rotational acceleration (Zhang’s study)—in order to deepen our understanding of the roles of neck structures in TBI mechanics. Our results demonstrated the impacts of both neck muscle activation and neck damping on HIC, BrIC, and brain MPS injury metrics across all simulated scenarios.

During head-neck impacts, the impact energy (forces) transmitted through the scalp, skull, and CSF to the brain can cause brain structural deformation. Neck muscles play a key role in attenuating this impact force. The level of this impact attenuation by neck muscles depends on their activation level, position, timing, and designated roles [11, 54]. For example, neck flexor muscles play a crucial role in counteracting the external impact forces in instances like the direct impact of a Golf ball on the forehead. On the other hand, in high acceleration scenarios like NBDL or Zhang’s helmet-to-helmet impacts, neck extensor muscles play a vital role in reducing neck hyper-flexion (in NBDL simulations), whereas neck rotator muscles contribute to reducing rapid neck rotational acceleration (in Zhang’s scenarios). The coordinated actions of these neck muscles also contribute to reducing the brain-skull relative motion. Consequently, across all NBDL and Golf ball impact simulations, our results exhibited a consistent pattern of reduced TBI metrics with the increase in neck muscle activation, which align well with prior experimental [20, 55] and FE-based computational studies [11, 17]. Furthermore, the increase in neck muscle activation resulted in reduced brain deformation in both frontal (coup) and occipital (countercoup) regions, suggesting that the strengthening of neck muscles through targeted training could be a crucial strategy in reducing the risk and severity of brain injuries in various impact scenarios. It also highlights the importance of the inclusion of neck muscles while investigating the TBI mechanism using both experimental and computational methods.

The brain selects the most appropriate muscles and varied activation levels (i.e., muscle activation strategies) to achieve a specific motor goal or movement. In Zhang’s linear and rotational acceleration scenarios, it was observed that increasing muscle activation from no-activation (5%) level to low-to-medium activation level (25%) resulted in a reduction in all brain injury metrics. However, when the muscle activation level was increased to the high muscle activation level (80%), there was a drastic increase in brain injury metrics. This inconsistent pattern revealed two unprecedented insights. First, a simple muscle activation strategy, which involved uniformly applying high activations (80%) across flexor, extensor, and rotator muscles, might result in abnormally high joint reaction forces to the skull, which, in turn, can cause increased brain-skull movement. A more sophisticated and accurate muscle activation strategy, such as modeling the rotator muscles with a higher activation than the flexor or extensor muscle groups, might have consistently provided reduced brain injury metrics. Therefore, it is essential to model neck muscles with scenario-specific precise muscle activation strategies in order to obtain an accurate understanding of TBI mechanism. Secondly, athletes with a history of concussion, who typically exhibit abnormal muscle activation strategies, are at a higher risk of experiencing recurring TBI or other musculoskeletal injuries [56]. Thus, our findings underscored the importance of neck muscle activation strategies (i.e., motor function) in TBI study.

In this study, we perturbed neck damping by altering the viscoelastic properties of intervertebral discs in order to investigate the resulting changes in the brain’s mechanical response. Like mechanical dampers, higher damping characteristics of the neck are expected to absorb more impact energy as it leads to a greater phase difference between impact force and tissue response instances. Previous research [12] showed that increasing neck damping in a direct head impact reduces the adverse mechanical responses of the brain. Our findings align with this, demonstrating that neck damping leads to a reduction in brain injury metrics. Therefore, it is essential to model intervertebral discs with appropriate damping characteristics in order for an accurate and comprehensive understanding of the TBI mechanism. Interestingly, the reductions in brain injury metrics were more prominently observed in brain MPS, followed by BrIC, than HIC in all simulated scenarios. This could be due to the fact that MPS directly reflects the mechanical deformation (strain) of brain tissues and BrIC accounts for rotational head kinematics, whereas HIC is limited to linear head kinematics [57]. Though the brain MPS is a more reliable measure of TBI, it requires computational brain models, thus posing practical challenges in many applications. In contrast, both HIC and BrIC are kinematics-based metrics. Especially, owing to its ease-of-use in estimating a range of TBI severities (AIS 1 to 5) and the ability to capture the brain’s response to rotational accelerations or shear forces, BrIC has recently become a popular injury metrics in many field (both engineering and clinical settings) applications.

This study has several limitations. Although we modeled three scenarios to study TBI mechanisms, it is essential to acknowledge that our study did not encompass all head impact types. Future investigations should explore a broader range of impacts, especially rear and side impact scenarios, to enhance our understanding of the neck’s role in TBI mechanisms. Additionally, we assessed TBI using three commonly used injury metrics. The inclusion of other off-the-shelf TBI injury metrics could have provided a more comprehensive TBI assessment. Moreover, we examined the neck damping characteristics by perturbing the damping properties of cervical intervertebral discs. This may not fully capture the total neck damping properties as cervical muscles, ligaments, tendons, and other soft tissues also contribute to neck damping. Thence, future research should account their combined contribution to TBI mitigation. Furthermore, we attempted only three simplistic muscle activation strategies by separating neck muscles into two groups (flexor and extensor muscles). This may not fully reflect the intricate motor activation strategies governed by the brain. Furthermore, it’s essential to recognize that muscle activation strategies and neck properties are influenced by various factors, including brain impairments, age-related changes, and sports-related injuries. These factors should be taken into account in future research to understand the TBI mechanics of people with a history of those changes or disorders. Finally, we model each intervertebral disc as a single solid body, neglecting the inherent composition of annulus fibrosis and nucleus pulposus with distinct mechanical properties.

Despite these limitations, our study reveals the crucial role played by neck muscles and neck damping in mitigating the severity of brain injury during traumatic head impacts. The knowledge gained from this study has practical applications. These include the development of tailored neck strength training protocols to enhance the mechanical capacity of neck muscles and motor strategies, as well as the design of protective equipment for those at higher risk of neck injuries, with the purpose of reducing the likelihood of TBI during mechanical impacts.

## Acknowledgments

This work was primarily supported by the National Science Foundation (**2239110**). We thank Felipe Santos, Leonardo Wei, and Gustavo Paulon for their assistance in data processing and finite element model development.

## Statements and Declarations

The authors have no competing interest to declare.

